# Identification of a novel small molecule facilitating HIV elimination by the “Shock-and-Kill” approach

**DOI:** 10.64898/2025.12.03.692014

**Authors:** Yuichiro Hara, Haruki Kitamura, Kouki Matsuda, Sayaka Sukegawa, Kosuke Tanimoto, Chieko Fujisaki, Kenji Maeda, Hiroaki Takeuchi

## Abstract

The development of antiretroviral therapy (ART) has significantly improved the prognosis of people living with HIV (PLWH). However, reservoirs of latent viruses represent a major barrier to achieving a complete cure of HIV infection. To overcome the burden of viral latency, the “Shock-and-Kill” strategy employs latency-reversing agents (LRAs) to reactivate the virus and thus render it susceptible to elimination. Existing LRAs are not optimal and there is an unmet need for improvement. By screening a small molecule library, we identified the novel LRA candidate 2-HSB (2-hydrido-2,2’-spirobi(1,3,2-benzodioxaphosphole)) which not only reactivates latent HIV provirus but also induces specific cytopathic effects in both monocytic and T lymphocytic latent HIV reservoirs. Thus, this novel compound represents a dual “shock and kill” agent, unlike known LRAs which are limited to the “shock”. The probable involvement of HIV-1 *tat* protein may account for its selectivity. Although both shock and kill effects seemed to be regulated by spleen tyrosine kinase, they were independent of each other, indicating that neither activation-induced cell death nor cell death-induced reactivation was occurring. Crucially, 2-HSB induced viral transcription in ART-suppressed PLWH samples *in vitro.* Our findings present 2-HSB as a promising dual-action compound for the “Shock and Kill” strategy, demonstrating efficacy in both transformed cell lines and clinically-relevant primary samples. These results raise the possibility of distinguishing the mode of action of the two effects, potentially leading to the development of a single agent causing both HIV-1 reactivation and elimination and contributing to the improvement of treatment.

## 1. Introduction

Human immunodeficiency virus type 1 (HIV-1) infects CD4^+^ cells, including helper T cells and macrophages, and causes acquired immunodeficiency syndrome (AIDS). Since the discovery of the anti-HIV activity of azidothymidine (AZT) (1), effort has been made to control HIV infection, which has led to the development of antiretroviral therapy (ART) (2). However, only a few patients have achieved complete eradication of HIV, with the “Berlin patient” being the most widely recognised case (3). Currently, ART cannot cure HIV infection but it decreases the viral load sufficiently to prevent the development of AIDS; this is typically called a functional cure (4). The latent HIV reservoirs, which maintain a very low level of HIV transcription, are the major obstacle preventing HIV elimination (5). To achieve the eradication of HIV from cell reservoirs and truly cure the infection, multiple approaches are being applied, such as gene editing by the CRISPR-Cas9 system (6) and vaccines to enhance HIV-1-specific cytotoxic T cells (7). The “Shock-and-Kill” strategy is a different approach to eliminating HIV reservoirs (8). As the name suggests, this strategy consists of two steps: reactivating the latent provirus (Shock) by latency-reversing agents (LRAs) and then eliminating reactivated HIV reservoirs (Kill) by ART and the host immune responses.

Several types of LRAs with different mechanisms of action have been developed thus far (9). These include histone modifiers, bromodomain inhibitors, protein kinase C agonists, and Toll-like receptor agonists. However, although these LRAs achieved good reactivation of HIV-1 from cell reservoirs *in vitro*, none has resulted in a significant reduction of reservoir size *in vivo* (10, 11). Thus, there is an unmet need to develop novel LRAs with a new mode of action, optimally both reactivating HIV and directly eliminating reservoirs.

Here, we report the identification of a novel LRA candidate through screening a small chemical library, as previously described (12). This agent both reactivated the HIV-1 provirus and also mediated cytopathic effects on the HIV-1 reservoir cells. Hence, it may be considered not so much as an LRA but as a dual “Shock”-and-“Kill” agent.

## 2. Results

### 2.1 2-hydrido-2,2’-spirobi(1,3,2-benzodioxaphosphole) reactivates HIV-1 in both a latent HIV-1 cell-line model and ART-suppressed patientś cells

We identified 2-hydrido-2,2’-spirobi(1,3,2-benzodioxaphosphole), here designated “2-HSB”, as a novel LRA candidate through small molecule library screening using THP-1 Nluc #225 cells (Fig. 1A). To further investigate its reactivation parameters, the T cell-derived reservoir cell line J-Lat 10.6 (10.6) as well as the monocytic line THP-1 Nluc #225 (#225) were treated with 2-fold serial dilutions of this compound. In the *nef* region of the HIV-1 provirus, #225 cells possess NanoLuc protein (13), and 10.6 cells possess EGFP (14), such that the level of expression of these reporter proteins indicates the reactivation of the provirus. Prior to testing reactivation, we calculated the 50% cytotoxicity concentration (CC50) of 2-HSB on these model cell lines by measuring the amount of intercellular ATP as an indicator of viability. The CC50 of 2-HSB on #225 and 10.6 cells was 5.222 and 7.571 μM, respectively (Fig. 1B). Next, HIV-1 reactivation was determined by measuring the activity of the NanoLuc and EGFP reporters. PEP005 (PKC agonist) (15) and JQ1 (BETi) (16) acted as controls. This revealed that 2-HSB increased the expression of reporter proteins in both #225 cells and 10.6 cells (Supplementary Fig. 1A, 1B) in a dose-dependent manner. To assess reactivation of the HIV-1 provirus at the level of transcription and exclude the non-specific effects of 2-HSB itself on the reporter proteins, total cell RNA from 2-HSB-treated #225 and 10.6 cells were extracted in order to conduct RT-qPCR of HIV-1 *tat* mRNA. Consistent with the results from the reporter protein measurements, it was confirmed that 2-HSB reactivated the HIV-1 provirus in both cell lines (Fig. 1C).

**Fig. 1.**
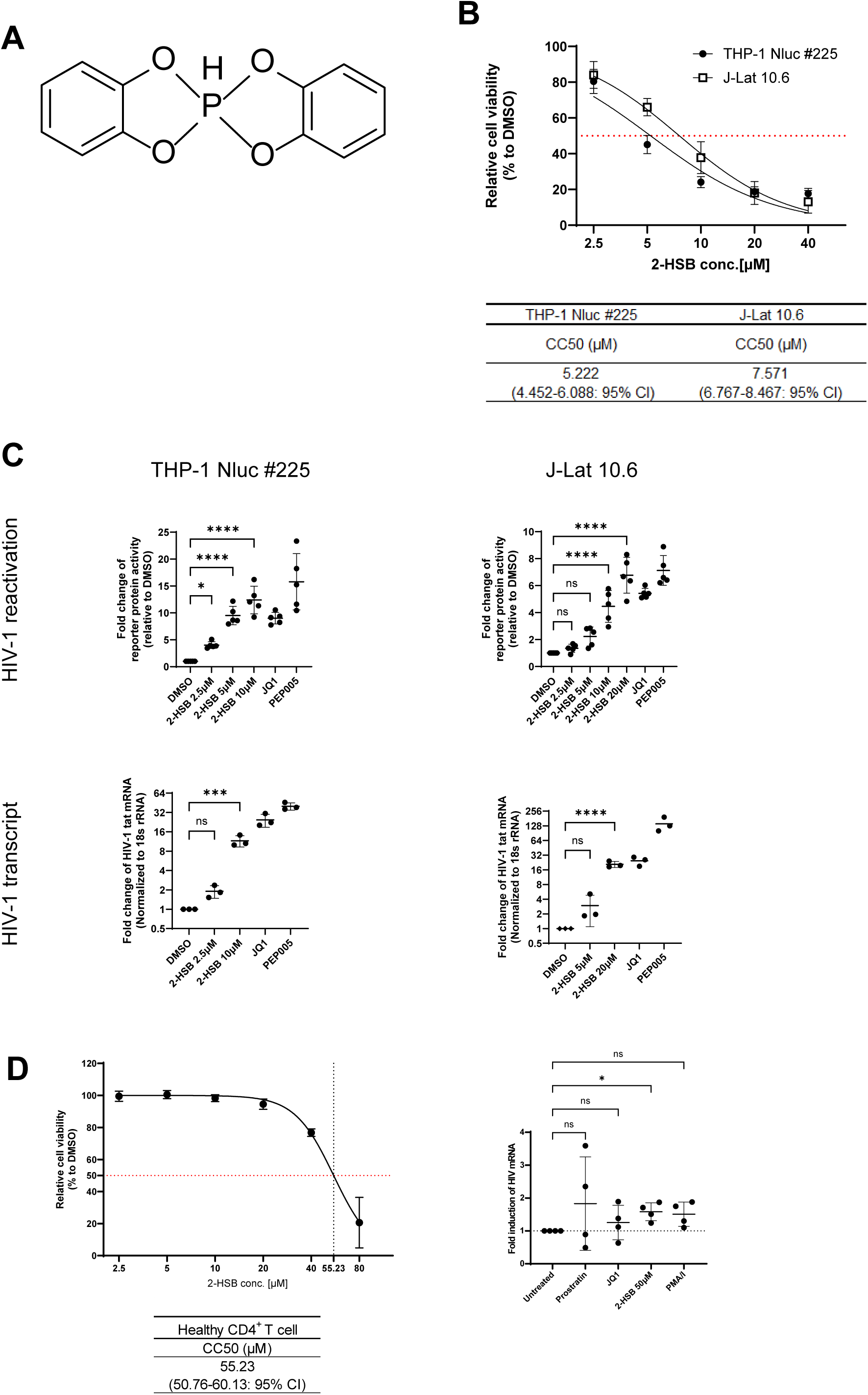
2-HSB reactivates latent HIV-1 in both a latent HIV-1 cell-line model and primary CD4^+^ T cells from ART-suppressed PLWH. (A) Chemical structure of 2-HSB. (B) Both #225 and 10.6 cells were treated for 24 hours with 2-fold serial dilutions of 2-HSB starting at 40 μM. The amount of intercellular ATP was measured as an indication of cell viability. Viability of DMSO-treated cells was set at 100% for calculating relative viability. CC50 values, the concentration required to reduce the viability of DMSO-treated cells by 50%, were calculated. (C) HIV-1 reactivation (upper panels) and HIV-1 transcripts (lower panels) following treatment with 2-HSB for 24 hours. #225 cells were lysed in the plates after treatment, and NanoLuc reporter activity was measured (upper left-hand panel). Reporter activity was normalised to intracellular ATP. The fold-change of normalised reporter activity relative to DMSO-treated cells is shown as the mean and SD calculated from five independent experiments. The number of GFP-positive 10.6 cells in surviving Via-Probe unstained cells was measured by FCM (upper right-hand panel). The fold- change of GFP-positive cells relative to DMSO-treated cells is shown as the mean and SD calculated from five independent experiments. Total cell RNA was extracted using RNeasy mini kits after treatment. HIV-1 transcripts (HIV-1 *tat* mRNA) were measured by RT-qPCR. Transcription level was calculated by the ΔΔCt methods using 18S ribosomal RNA as an internal reference. The means and SD calculated from three independent experiments are shown (lower panels). (D) 2-HSB effectively reactivates latent reservoirs from ART-suppressed patients *in vitro.* (Left) Primary CD4^+^ T cells isolated from three independent healthy donors were treated for 24 hours with 2-fold serial dilutions of 2-HSB starting at 80 μM, and CC50 was determined as in Fig. 1B. (Right) Primary CD4^+^ T cells isolated from ART-suppressed patients were treated for 24 hours with or without LRAs. Total cell RNA was collected after the treatment, and the amount of HIV-1 mRNA was measured by RT-qPCR. The fold-induction relative to untreated samples is plotted. The means and SD calculated from four independent samples are shown. Statistical significance was determined by one-way ANOVA and Tukey’s post-hoc multiple comparison testing. ns., not significant (*P*>0.05), * *P*<0.05, **** *P*<0.0001.

Validating the efficacy of LRAs in primary cell models is essential for clinical translation. We first assessed the safety of 2-HSB for healthy primary CD4^+^ T cells. As shown in Figure 1D, the CC50 value of 2-HSB was found to be 55.23 μM (Fig. 1D, left panel). Based on the CC50, the test concentration of 2-HSB on primary CD4^+^ T cells from ART-suppressed patients was set at 50 μM. While the reactivation of latent HIV by conventional LRAs (Prostratin and JQ1) failed in some patientś samples, PMA / Ionomycin combination and 50 μM 2-HSB reactivated HIV in all samples (Fig. 1D, right panel). Especially, 50 μM 2-HSB alone mediated statistically significant induction relative to the untreated control, suggesting its strong potential compared with established LRAs.

### 2.2 2-HSB selectively induces Caspase-3-dependent apoptosis in HIV-1 reservoir cell lines and depletes these cells from the population

As the term LRA refers only to the “Shock” part of the Shock-and-Kill strategy, it is often pointed out that the “Kill” must not be ignored (17, 18). Because the ultimate aim of the strategy is to eradicate HIV-1-infected cells residing in people living with HIV (PLWH), it is necessary to consider how to eliminate the reservoir cells. As we confirmed by the data shown in Fig. 1B, 2-HSB exerted a cytopathic effect on latent reservoir cells. To investigate whether this cytopathic effect was selective for such cells, we conducted the same experiment as in Fig. 1B on heterogeneous populations derived from mixing #225 or 10.6 cells with their parental cells, THP-1 and Jurkat. Encouragingly, 2-HSB had little effect on the parental cells (CC50 > 40 μM for both THP-1 and Jurkat cells). This implies that the cytopathic effect requires the presence of the HIV-1 provirus. This preferential induction of cell death was thus conserved in a heterogeneous population of HIV-1-positive and negative cells, as the percentage of HIV-1-positive cells and the level of cytopathic effect showed a positive correlation (Fig. 2A). These results indicated that 2-HSB caused cells death specifically in HIV-1-positive cells without causing the bystander cell death of HIV-1-negative cells despite the significantly enhanced HIV-1 provirus activity at the same time.

**Fig. 2.**
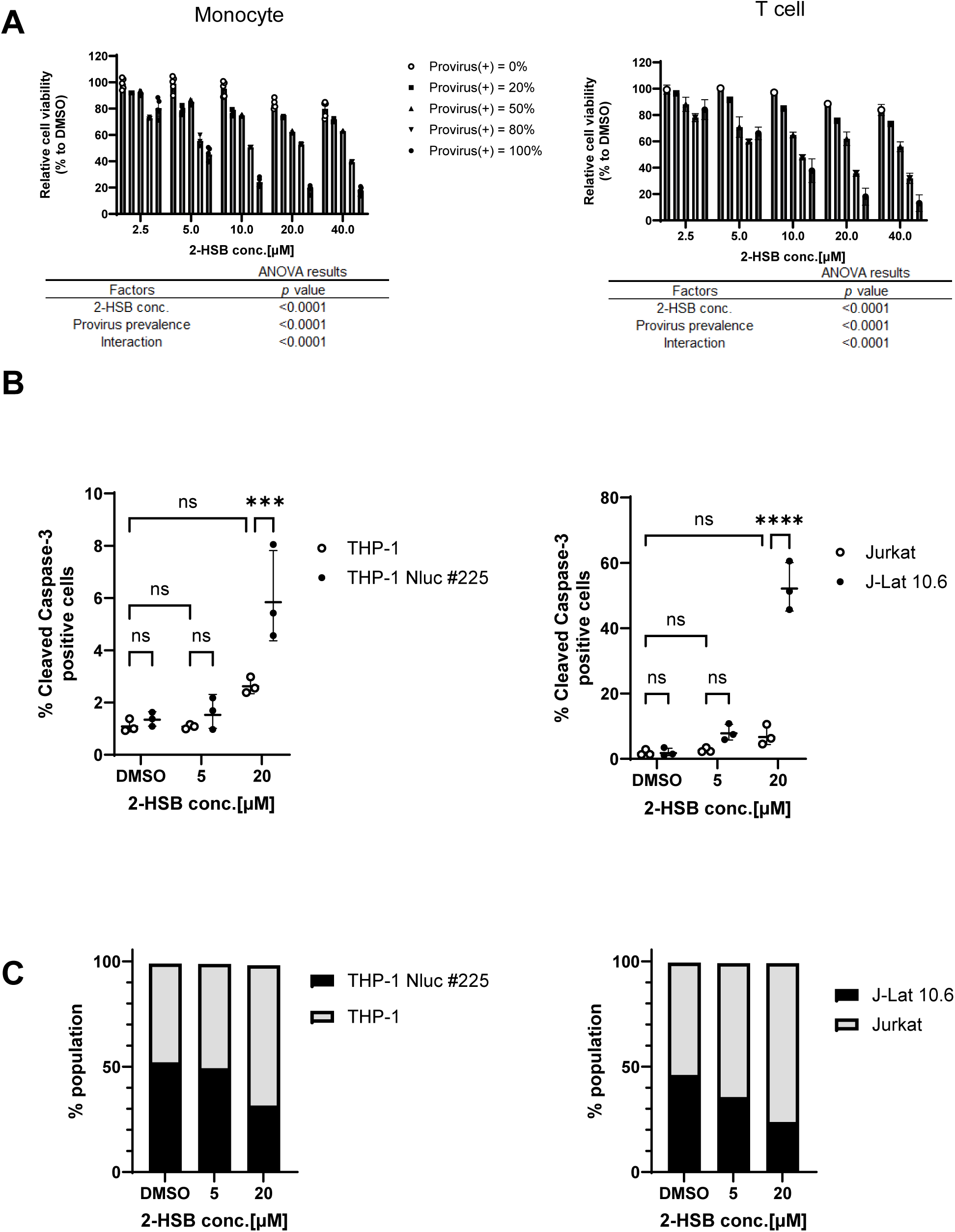
Selective induction of Caspase-3-dependent apoptosis on the HIV-1 reservoir. (A) THP-1 and #225 cells (left), Jurkat and 10.6 cells (right) were mixed to create a variable percentage of HIV-1 provirus-positive cells in the population, and treated with 2-fold serial dilutions of 2-HSB for 24 hours. The amount of intercellular ATP was measured to indicate cell viability. Viability of DMSO-treated cells was set at 100% for calculating relative viability. The results are shown as mean values and standard deviation calculated from at least three independent experiments. The effects of the presence of the provirus and of 2-HSB were determined by two-way ANOVA. (B-C) CellTracker-labeled HIV-1-negative cells (THP-1 or Jurkat) were mixed 1:1 with unlabeled HIV-1-positive cells (#225 or 10.6) and treated with 2-HSB for 24 hours. Apoptosis markers and population dynamics were analyzed by flow cytometry. (B) The percentage of cleaved Caspase-3-positive cells was quantified within the labeled (parental) and unlabeled (HIV-1-positive) populations. Left: Monocytic lineage (THP-1 vs. #225). Right: T-lymphocytic lineage (Jurkat vs. 10.6). (C) Specific depletion of HIV-1-positive cells. The population distribution (% of total cells) of labeled and unlabeled cells following treatment. Left: Monocytic lineage. Right: T-lymphocytic lineage. The means and SD calculated from three independent experiments are shown. Statistical significance was determined by two-way ANOVA. ns., not significant (*P*>0.05), *** *P*<0.001, **** *P*<0.0001.

To rigorously evaluate the selectivity of 2-HSB-induced apoptosis in mixed population, we measured the levels of cleaved Caspase-3, which is known to be involved in the apoptotic pathway (19). THP-1 or Jurkat cells were labelled with a fluorescent probe and mixed with unlabelled #225 or 10.6 cells. The mixed population was then treated with 2-HSB for 24 hours, and the amount of cleaved Caspase-3 was measured in unlabelled/labelled cells by flow cytometry. Similar to the previous results, 2-HSB treatment resulted in a significant induction of cleaved Caspase-3 exclusively within the HIV-1-positive #225 or 10.6 cell population, while no significant increase in apoptosis was observed in the HIV-1-negative THP-1 or Jurkat population (Fig. 2B and Supplementary Fig. 2). Furthermore, this selective killing led to a specific depletion of HIV-1-positive cells from the co-culture over time (Fig. 2C and Supplementary Table 1). These results conclusively demonstrate that 2-HSB targets HIV-1-positive cells for elimination.

### 2.3 SYK tyrosine kinase regulates the effects of 2-HSB

NF-κB is a key gene involved in both the reactivation of HIV-1 and apoptosis of infected cells (20). Therefore, we sought genes involved in the NF-κB signalling pathway that were upregulated only in provirus-positive cells upon exposure to 2-HSB. Using RNA microarray analysis, we found that the transcription of SYK which is involved in NK-κB signalling pathway was markedly upregulated only in HIV-1 provirus-positive cells (Supplementary Fig. 3 and Supplementary Table 2). In addition to its pro-apoptotic effect on cells (21, 22), it is known to regulate the HIV-1 reactivation by some LRAs, including PKC agonists and TLR8 agonists (23). To determine whether SYK was also a mediator of the effects of 2-HSB, we applied a commercially available SYK inhibitor, SYK inhibitor II dihydrochloride (SYK-IN-II) (24). Cells were pretreated with different concentrations of SYK-IN-II for 24 hours and then treated with 2-HSB to test the cytopathic effects and HIV-1 reactivation. We found that inhibition of SYK significantly reduced the cytotoxicity of 2-HSB in a dose-dependent manner (Fig. 3A) as well as HIV-1 reactivation in both #225 and 10.6 cells (Fig. 3B). These results indicated that SYK is an indispensable cellular factor for 2-HSB-mediated cytotoxicity and HIV-1 reactivation in both monocyte- and T cell-derived HIV-1 provirus-positive cells. From these results, we concluded that 2-HSB was capable of reactivating the HIV-1 provirus and eliminating provirus-positive cells in both monocyte and T cell reservoir populations via an identical pathway. The finding that 2-HSB targeted not only T cells but also monocytes is encouraging for its application in the clinic, as there is a long-standing awareness of T cell reservoirs (25) but nowadays also an increasing concern about non-T cell reservoirs(26).

**Fig. 3.**
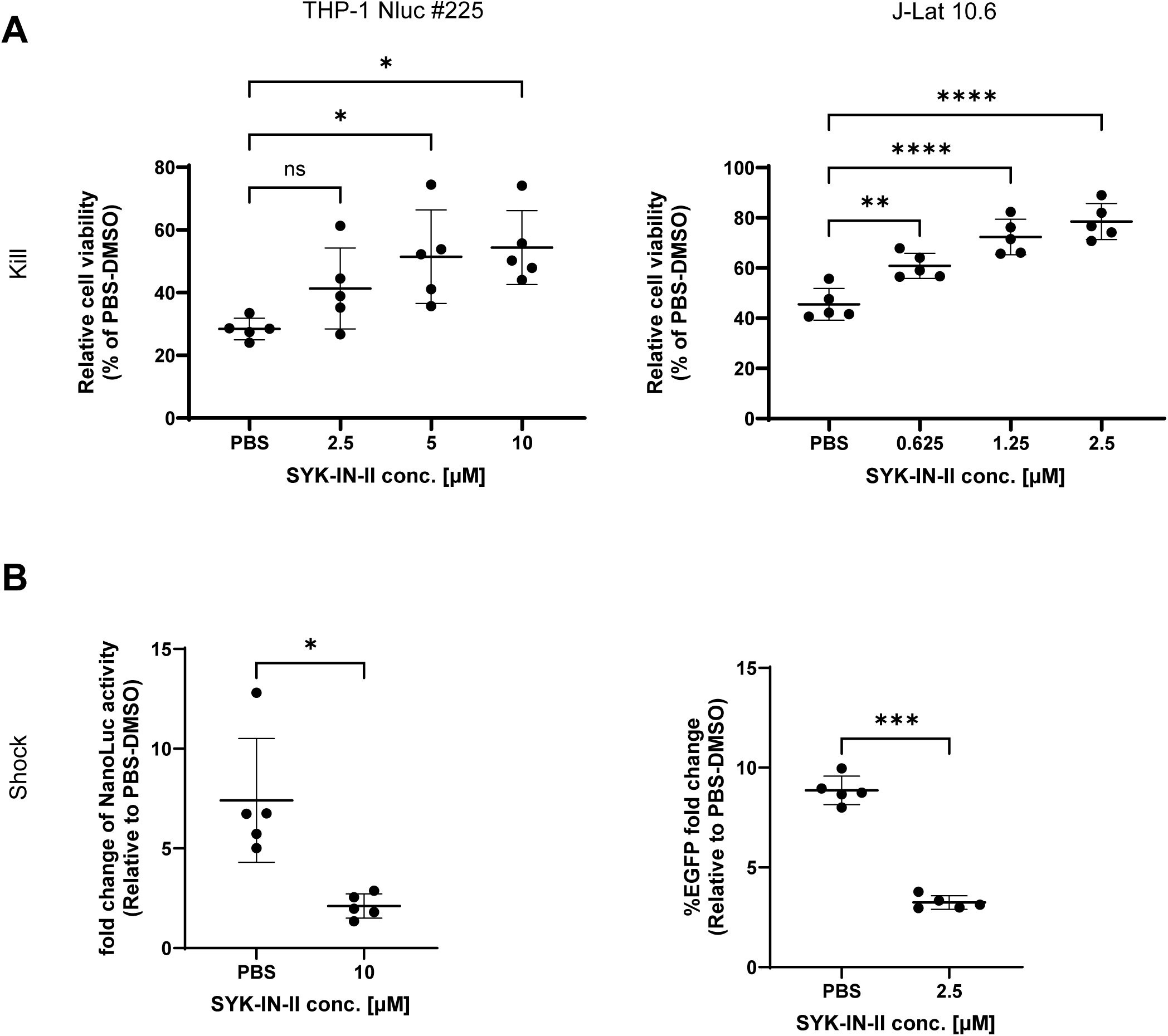
SYK spleen tyrosine kinase is an indispensable host factor for the action of 2-HSB. (A, B) #225 and 10.6 cells were pretreated with SYK-IN-II for 24 hours and then cells were treated with 2-HSB for 24 hours. Cell viability (Kill) and HIV-1 reactivation (Shock) were measured as in Fig. 1B and Fig. 1C. The means and SD calculated from five independent experiments are shown. Statistical significance was determined by one-way ANOVA and Tukey’s post-hoc multiple comparison testing or two-tailed Student’s t-testing. ns., not significant (*P*>0.05), * *P*<0.05, ** *P*<0.01, *** *P*<0.001, **** *P*<0.0001.

### 2.4 HIV-1 Tat protein is a crucial viral factor for the action of 2-HSB

Because it is well-established that several different HIV-1 viral proteins are involved in maintaining latency (27) and causing cell death (28, 29), we investigated the involvement of these viral proteins in the effects of 2-HSB. Due to its dual effects, HIV-1 reactivation and selective HIV-1 provirus-positive cells killing, we hypothesized that HIV-1 Tat was a possible viral factor targeted by 2-HSB, because it is a crucial protein for both HIV-1 transcription and HIV-1-induced cell death (30). We compared the effects of 2-HSB on J-Lat A2 and J-Lat A72 cells to determine the importance of HIV-1 Tat. Like 10.6 cells, both of these cell lines are derived from Jurkat, but they do not harbour the full-length HIV-1 provirus. However, they do contain a partial-length provirus, one with and one without the *tat* gene (Fig. 4A). We conducted reactivation and cytotoxicity assays in the same manner as with 10.6 cells. While 2-HSB mediated significant reactivation in a dose-dependent manner in A2 cells, it had no effect in A72 cells (Fig. 4B). This pattern was reproduced with the cytopathic effect as well in that A2 and 10.6 cells were similarly susceptible, but A72 cells were more resistant albeit the effect was still significantly stronger than that in Jurkat, the HIV-1 provirus-negative parental control cells (Fig. 4C). These results suggest that the HIV-1 Tat protein is an indispensable viral factor required for the effects of 2-HSB. The finding that the actions of 2-HSB are dependent on HIV viral factor strengthens the case for its potential clinical application, as there would be no negative side-effects on uninfected tissues.

**Fig. 4.**
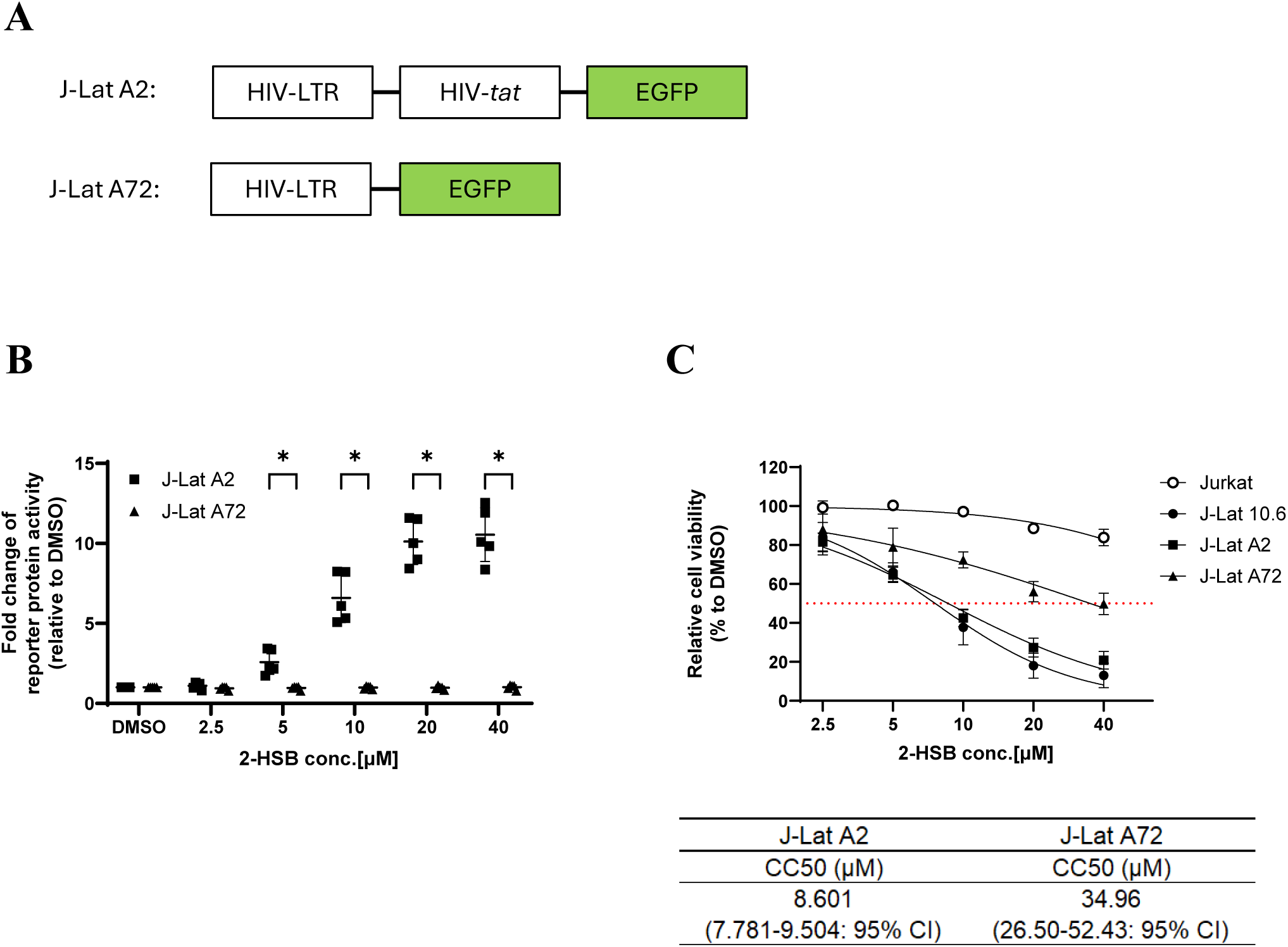
The effects of 2-HSB are HIV-1 Tat-dependent. (A) Schematic diagrams of HIV-1 provirus in A2 and A72 cells. (B) Reactivation of HIV-1 in A2 and A72 cells was measured following 24 hours of treatment with 2-HSB. The number of GFP-positive cells in surviving Via-Probe-unstained cells was measured by FCM. The fold- change of GFP-positive cells relative to DMSO-treated cells is shown as the mean and SD calculated from five independent experiments. (C) A2 and A72 cells were treated for 24 hours with 2-fold serial dilutions of 2-HSB starting at 40 μM. The amount of intercellular ATP was measured to indicate cell viability. Viability of DMSO-treated cells was set at 100% for calculating relative viability. CC50, the concentration required to reduce the viability of DMSO-treated cells by 50%, was then determined. Results of Jurkat (Fig. 2A) and 10.6 cells (Fig. 1B) are also depicted. The means and SD calculated from five independent experiments are shown. Statistical significance was determined by two-tailed multiple t-test. * *P*<0.05.

### 2.5 Transcriptomic analysis of cells exposed to 2-HSB reveals enrichment of the apoptosis-signalling pathway

Certain known LRAs, such as HDAC inhibitors, BET bromodomain inhibitors (BETi) and PKC agonists, reactivate the HIV-1 provirus by manipulating the host cell transcription dynamics (16, 31, 32). Accordingly, we conducted transcriptomic analysis to confirm whether 2-HSB also altered the transcriptome dynamics. Gene Ontology (GO) enrichment analysis of significantly upregulated genes showed that the term apoptotic process had the highest enrichment scores in both #225 and 10.6 cells (Fig. 5A), supporting the pro-apoptotic effects of 2-HSB at the transcriptional level. To explore any similarities and differences between 2-HSB and typical LRAs, enriched GO terms for genes significantly upregulated following JQ1 (BETi) and Prostratin (PKC agonist) (33) treatments were compared with that of 2-HSB (Fig. 5B). This revealed that 2-HSB behaves as a broad activator similar to Prostratin, although the results suggested that it is functionally more similar to JQ1 than to Prostratin (2-HSB and JQ1 = 12 out of 22 GO terms [54.5%], 2-HSB and Prostratin 83 out of 218 GO terms [38.1%]). Because JQ1 is reported to reactivate HIV by a tat-dependent mechanism (16), these results are consistent with the possible tat-dependency of the action of 2-HSB (Fig. 4B and 4C).

**Fig. 5.**
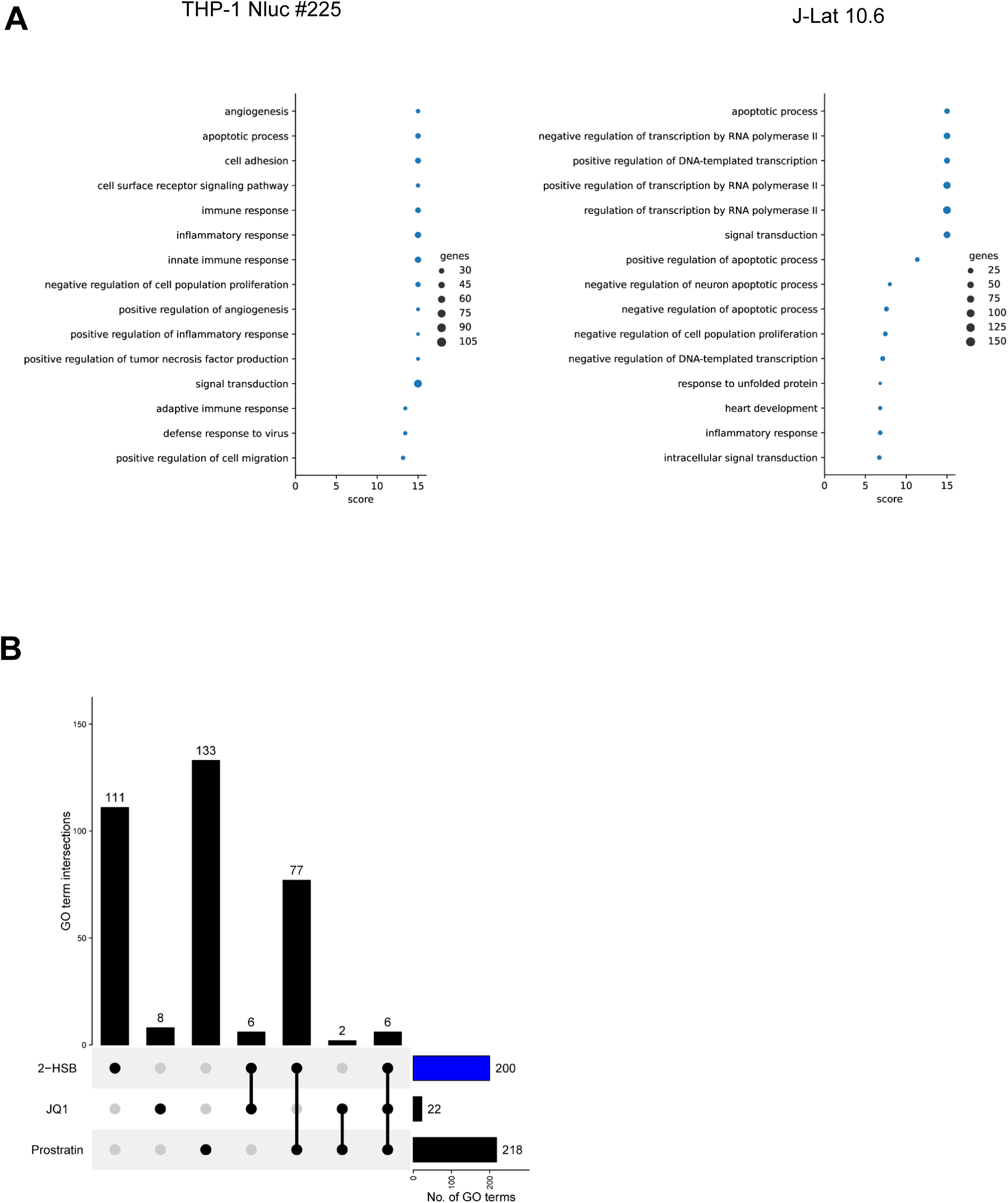
Transcriptomic analysis reveals upregulation of apoptotic processes in HIV reservoir cells treated with 2-HSB. (A) GO enrichment analysis using RNA-seq data of treated #225 (right) and 10.6 (left) cells. The GO terms of the significantly upregulated gene sets (FDR < 0.05, log2 fold change > 1) were extracted. (B) Upset plot of intersections between significantly enriched GO terms under different treatments. GO terms that were enriched in both #225 and 10.6 cells were used for comparison. The bar chart on the right shows the total number of enriched GO terms under each treatment. The bar chart on the top indicates the number of enriched GO terms at each intersection.

### 2.6 Only one or other of the dual effects of 2-HSB is seen in HIV-1 reservoir cells

Since activated HIV-1 may cause apoptosis (34) and reciprocally the induction of apoptosis may reactivate latent viral reservoirs (35), we elected to investigate whether the dual *Shock* and *Kill* effects of 2-HSB were sequential or simultaneous. Briefly, 10.6 cells were stimulated with a high concentration of 2-HSB for 24 hours, followed by staining of cleaved Caspase-3 as an indicator of apoptosis induction. The cells were analysed by flow cytometry to determine whether 2-HSB was causing only one outcome or both simultaneously at the individual cells (Fig. 6A). We compared these features with JQ1 (Fig. 6B), for which the CC50 reported previously (9.51µM, (12)) was similar to 2-HSB (7.571µM, Fig. 1B). The proportions of three populations were compared, namely, only EGFP-positive (Shock only), only cleaved Caspase-3-positive (Kill only), and both EGFP and cleaved Caspase-3-positive (Shock & Kill). While there were no differences between 2-HSB and JQ1 on the properties of cells indicating only Shock, the percentages of Kill-only and Shock & Kill-double positive cells were significantly higher with JQ1 than with 2-HSB (Fig. 6C). Furthermore, the percentage of Shock & Kill-double positive cells within the cleaved Caspase-3-positive population was significantly lower with 2-HSB; in fact, there was no significant difference from the DMSO control (Fig. 6D). This indicates that 2-HSB caused neither HIV-1-induced cell death nor cell damage-induced activation, or a “random” cytopathic effect. Besides, 2-HSB induced either only reactivation or cell death in individual cells, suggesting the existence of precise cellular regulators of these two effects.

**Fig. 6.**
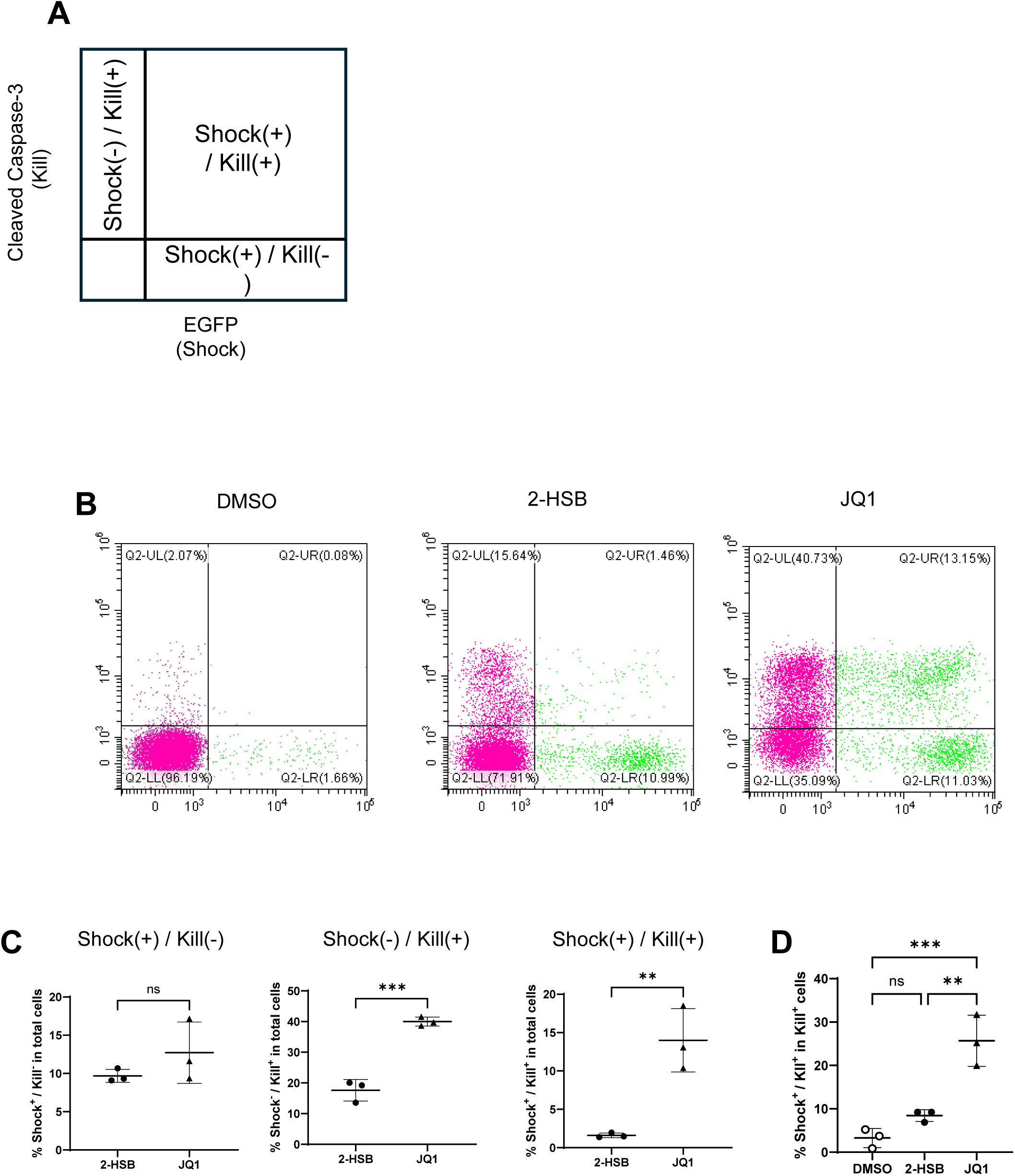
The *Shock* and *Kill* effects of 2-HSB are independent of each other. (A) Schematic diagrams of flow cytometry analysis. (B) 10.6 cells were treated for 24 hours with 2-HSB (left) or JQ1 (right) at 20 μM followed by staining for intercellular cleaved Caspase-3 using Cleaved Caspase-3 Staining Kits (red); the number of cleaved Caspase-3/EGFP-double-positive cells was measured by FCM. (C) The percentages of cells in Shock(+) (left) / Kill(-), Shock(-) (middle) / Kill(+) and Shock(+) / Kill(+) (right) fractions within the total cell population. (D) The percentage of Shock(+) / Kill(+) cells within the fraction of Kill(+) cells. The means and SD calculated from three independent experiments are shown. Statistical significance was determined by two-tailed Student’s t-testing (C) or one-way ANOVA (D). ns., not significant (*P*>0.05), * *P*<0.05, ** *P*<0.01, *** *P*<0.001.

## 3. Discussion

In attempts to reactivate latent HIV from reservoirs impermeable to ART and to render the virus susceptible to destruction, multiple LRAs with different modes of action have been developed (9). However, their promising *in vitro* reactivation capabilities are not translating to the success of clinical trials aimed at eradicating HIV-1 reservoirs in patients (10, 11). Thus, there is an unmet need to explore and develop new LRAs with novel modes of action. Here, we identified and evaluated a novel LRA candidate, designated 2-HSB, which serves as a *Shock “AND” Kill* agent at the cell population level (Fig. 7A) while it acts as a *Shock “OR” Kill* agent at the single cell level (Fig. 7B). This compound is capable of both reactivating HIV in latent reservoir cells and also inducing their selective apoptosis without damaging uninfected cells of the same lineage, resulting in their selective depletion from a mixed population (Fig. 2B, Fig. 2C). Notably, 2-HSB acted on both monocyte- and T cell-derived populations in identical pathways (Fig. 5A). In both lineages, SYK was found to act as one of the key host factors required for the activity of 2-HSB (Fig. 3A, Fig. 3B). Currently, T cells are considered to be the most important HIV-1 reservoir populations (25, 36). More recently, a role of monocytic cell reservoirs has been increasingly implicated (37), and means to eradicate all types of reservoirs are being intensively investigated (38). Thus, the findings that 2-HSB mediates similar effects in both types of reservoir populations with an identical mode of action presented here encourage continuous future studies on its use in the clinic.

**Fig. 7.**
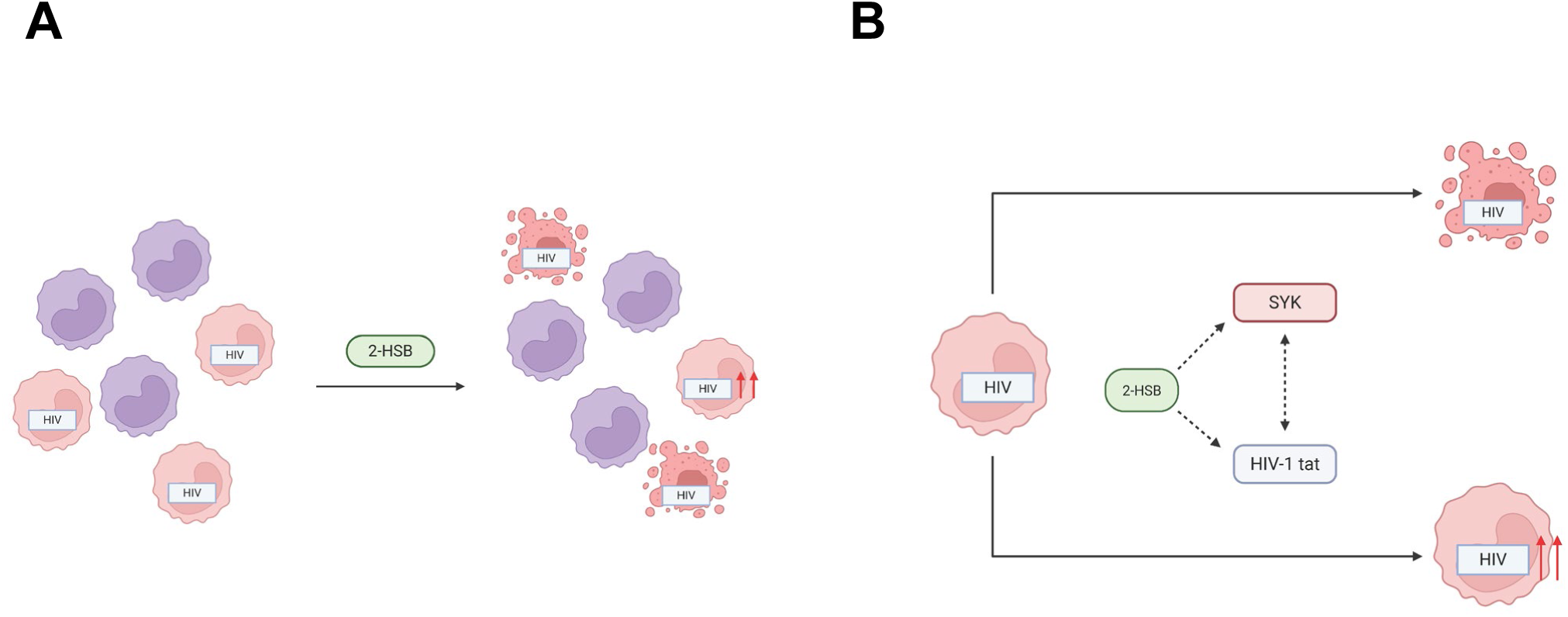
Proposed model of the 2-HSB mechanism of action targeting HIV-1 latent reservoirs. (A) Population-level effect (“Shock AND Kill”). Within the total cell population, 2-HSB functions as a dual-action agent, inducing concomitant viral reactivation and cell death. (B) Single-cell fate (“Shock OR Kill”). At the single-cell resolution, 2-HSB triggers mutually exclusive outcomes; individual latently infected cells undergo either viral reactivation (“Shock”) or apoptosis (“Kill”), but not both concurrently. Figures were created with BioRender.com (Hara, Y. 2025).

The HIV-1 Tat protein is a crucial viral factor for the biology of HIV-1, not only being essential for efficient HIV-1 transcription and replication (39), but also for manipulating host gene expressions to create favourable HIV environments (40). As it is a viral protein, targeting HIV-1 Tat as a therapeutic strategy might offer increased safety of treatments due to virus selectivity. The finding that 2-HSB essentially failed to induce Caspase-3-dependent apoptosis of HIV-1 provirus-negative cells (Fig. 2A, Fig. 2B) reinforces this notion of safety, as its effects seemed to be Tat-dependent. This is illustrated by our findings that its abilities to reactivate HIV and to induce cell death were almost absent (Fig. 4B) and nearly halved (Fig. 4C) in Tat-negative model cells, respectively. These results are also supported by transcriptomic analysis, which suggested that the mechanism of action of 2-HSB was similar to JQ1, that is thought to be Tat-dependant (Fig. 5B). While HIV reactivation by 2-HSB was heavily dependent on Tat, the HIV-1 long terminal repeat (LTR) itself also played a major role in its cytopathic effects. As it is well-known that Tat*-*defective provirus can nonetheless produce viral progeny (41), even though Tat dysfunction might lead to latency (42), reservoirs with Tat-defectives are also an important target. Thus, understanding how 2-HSB induced cell death in Tat-defective cells, albeit at a modest level, is worth investigating. While the involvement of different viral proteins in the apoptosis of HIV-1-infected cells is well-studied (43), little is known about the pure role of the HIV-1 LTR alone for apoptosis. As the provirus of A72 cells, which we used to demonstrate the effect of Tat-deficits, consists of only HIV-1 LTR and EGFP (Fig. 4A), our results imply a novel role of HIV-1 LTR itself for the pathogenicity of HIV-1 infection, although these findings require further clarification. Collectively, we have identified both host and viral factors involved in 2-HSB activity though their precise functional relationship remains to be determined.

Interestingly, despite the reactivation and cytopathic effects of 2-HSB being dependent on the same host and viral factors, these two effects were found to be independent of each other. We showed that even at high concentrations of 2-HSB, which killed >80% of HIV-1 reservoir cells, those cells in which HIV-1 reactivation was induced and those in which Caspase-3-dependent apoptosis was observed were clearly separated. Moreover, the percentage of cells with HIV-1 reactivation within the apoptotic population was not different from that of the control (Fig. 6B). This feature has two strong merits when considering the clinical use of 2-HSB. First, it would be able to deplete most of the reservoirs without causing excess inflammation. It is clear that active HIV and apoptosis may both induce strong inflammatory responses (44, 45), and there is little doubt that a combination of both would make the inflammation worse. Thus, if we aim to eradicate the HIV reservoir cells, it is desirable to remove them without stimulating untoward inflammatory responses. Accordingly, the cytopathic effect induced by 2-HSB would be ideal because it does not cause HIV reactivation in dying cells. Second, we may still be able to achieve a high level of HIV reactivation, because 2-HSB does not induce apoptosis of those cells with reactivated virus. This may seem paradoxical, given the first advantage described above, because a high activation rate will cause inflammation, but it is a question of degree. Since the basis of the Shock-and-Kill approach is to reactivate HIV from the reservoirs and then eradicate the virus and the infected cells by combining ART and triggering host immune responses, it is necessary that viral production is maintained to induce an appropriate immune response. Although we do not yet know whether reservoirs reactivated by 2-HSB will maintain virus production persistently, at least they will not be eliminated by apoptosis. There would therefore be more opportunity to induce a sufficient immune response. Thus, the favourable characteristics of 2-HSB are the ability to decrease the size of the HIV-1 reservoir with minimal inflammation and to sustain HIV production without killing the reservoir cells.

Importantly, we have demonstrated the reactivation ability of 2-HSB on samples from ART-suppressed PLWH, which supports its application in clinical settings (Fig. 1D). Compared to model cell lines such as #225 and 10.6 cells, which are clonal populations, patients’ HIV-1 reservoirs are much more heterogeneous in several respects, such as integration sites (46, 47) and epigenetic modifications (48). Hence, demonstrating HIV-1 reactivation ability on different clinical samples is a critical supporting step for future studies.

Overall, we have documented a promising ability of 2-HSB to act as an LRA on both monocyte-derived as well as T cell-derived reservoirs, and *ex vivo* patient-derived clinical samples. However, the mode of action of 2-HSB is still not fully understood. Considering its use in the clinic, further investigation is necessary to prevent unexpected side effects and enhance safety. An additional limitation is that, due to the limited amount of available samples from ART-suppressed patients, we were unable to confirm 2-HSB’s cytotoxicity *in vitro*. To overcome these limitations, we are now conducting comprehensive omics analysis to identify key host factors. Furthermore, we are establishing an *in vitro* primary latent HIV-1 infection model to investigate the killing activity and host determinants of 2-HSB on clinically relevant samples. Nevertheless, it is already clear that our findings have provided new insights into possible Shock and Kill strategies and should contribute to the establishment of better treatments for PLWH.

## 4. Materials and Methods

### 4.1 Cell lines and cell culture reagents

THP-1, Jurkat, THP-1 Nluc #225 cells (#225) (13), J-Lat 10.6 cells (10.6), J-Lat A2 cells (A2) and J-Lat A72 cells (A72) (14) were cultured in RPMI-1640 (Nacalai Tesque) supplemented with 10% heat-inactivated FCS (Sigma Aldrich) and Penicillin/Streptomycin (Nacalai Tesque).

### 4.2 Chemical compounds

2-hydrido-2,2’-spirobi(1,3,2-benzodioxaphosphole) was obtained from Princeton BioMolecular Research. SYK Inhibitor II dihydrochloride was obtained from TargetMol. (+)-JQ1 (BRD4 inhibitor) was obtained from Adooq Bioscience. PEP-005 (PKC activator, ingenol-3-angelate) was obtained from Cayman Chemical Company.

### 4.3 Small molecule library screening

Screening was conducted as previously described (12). Briefly, 2 x 10^4^ of #225 cells/well were seeded into 96-well plates together with 10 μM of compounds for 24 hours. Nano-Glo Luciferase Assay System (Promega) was then used according to the manufacturer’s protocol to determine the NanoLuc activity. Compounds that caused more than 2 times higher induction of NanoLuc activity compared to unstimulated control were used for further analysis. NanoLuc activity was measured using the GloMax Multi-detection system (Promega).

### 4.4 Measurement of cell viability

A series of 2-fold dilutions of the test compounds was used to measure cell viability. Briefly, 2 x 10^4^ cells/well were seeded into 96-well plates together with the compounds for 24 hours. Viral ToxGlo (Promega) was then used according to the manufacturer’s protocol to determine the cellular ATP content using GloMax Multi-detection system (Promega). The CC50 was calculated by GraphPad Prism v10 using the amount of cellular ATP to indicate cell viability.

### 4.5 Reporter protein analysis

For #225 cells, 2 x 10^4^ cells/well were seeded into 96-well plates together with the compounds for 24 hours. NanoLuc activity was determined as described above. For 10.6, A2 and A72 cells, flow cytometry (FCM) analysis was conducted on a CytoFLEX SRT (Beckman Coulter). First, stimulated cells were washed with PBS(-) and stained with BD Via-Probe™ Cell Viability Solution (BD Biosciences) according to the manufacturer’s protocol. Cells were then fixed with 0.8% paraformaldehyde (PFA). Forward and side scatter parameters were used to distinguish viable cells within the total sample. Fluorescence channels PC5.5 and FITC were used to detect signals from Via-Probe and EGFP, respectively.

### 4.6 Quantitative RT-PCR

Total cell RNA was extracted using RNeasy mini kits (QIAGEN). Quantification of the doubly spliced form of HIV-1 NL_4-3_ associated RNA by quantitative RT-PCR (RT-qPCR) was performed as previously described (12). Verso SYBR Green 1-Step qRT-PCR ROX Mix (Thermo Scientific) was used to evaluate the amount of spleen tyrosine kinase (SYK) mRNA. Primer pairs 5’-CATGGAAAAATCTCTCGGGAAGA-3’ and 5’- GTCGATGCGATAGTGCAGCA-3’ were used to amplify and detect SYK mRNA. Amplification levels were normalised to GAPDH. All RT-qPCR was performed using the StepOnePlus Real-Time PCR system (Applied Biosystems).

### 4.7 Live cell labelling

Jurkat cells were collected and resuspended in fresh RPMI-1640 supplemented with 10% heat-inactivated FCS and Penicillin/Streptomycin at 1 x 10^6^ cells/mL. Cell Tracker Deep Red dye (Invitrogen) was added to the cells at a ratio of 1:1000 and incubated for 30 minutes. Cells were then washed twice with PBS(-) and resuspended in fresh RPMI-1640 supplemented with 10% heat-inactivated FCS and Penicillin/Streptomycin. The APC channel was used to detect signals from CellTracker Deep Red-stained cells by FCM.

### 4.8 Cleaved Caspase-3 analysis

1 x 10^6^ cells/well were seeded into 24-well plates together with the compounds for 24 hours. Cleaved Caspase-3 was then stained using Cleaved Caspase-3 Staining Kit (Red) (Abcam) according to the manufacturer’s protocol. After staining, cells were fixed with 0.8% PFA and the relative number of cleaved Caspase-3-positive cells was determined by FCM. The PE channel was used to detect signals from stained cleaved Caspase-3 by FCM.

### 4.9 Transcriptomic analysis

#225 and 10.6 cells were treated with compounds for 24 hours. Thereafter, dead and damaged cells were removed using Dead Cell Removal Kits (Miltenyi Biotech), and total RNA was extracted from viable cells. For this, the RNase-Free DNase Set (QIAGEN) was used to remove residual DNA from the sample processed by RNeasy mini kits (QIAGEN). RNA-seq was performed by TAKARA Bio, and fastq data were obtained for further analysis. Read data were mapped to human genome reference GRCh38 using STAR (49), and the transcription level was estimated by RSEM (50). The “gencode.v46.basic.annotation.gtf” file downloaded from Gencode was used for gene annotation. Differentially expressed genes (DEGs) were analysed by DESeq2(51). For GO enrichment analysis, GO terms for each gene were obtained using the “gene2go” file and the “gene2refseq” file downloaded from NCBI. GO enrichment was tested using hypergeometric distribution, and obtained p-values were corrected using the Benjamini-Hochberg (BH) method. Upset plots of enriched GO terms was made using ComplexHeatmap (52). RNA microarray analysis was performed by TaKaRa Bio Inc. (Otsu, Japan).

### 4.10 Protein inhibitor analysis

#225 and 10.6 cells were pretreated with SYK Inhibitor II dihydrochloride (TargetMol) for 24 hours and then treated with compounds for another 24 hours. Reporter protein activity and cell viability were measured as described above.

### 4.11 Measurement of cytotoxicity to primary CD4^+^ T-cells from healthy donors

Whole peripheral blood mononuclear cells (PBMC) were isolated from healthy donors’ blood by density-gradient centrifugation using Ficoll-Paque PLUS (cytiva), and CD4^+^ T-cells were purified using the human CD4^+^ T cell Isolation Kit (Miltenyi Biotec) according to the manufacturer’s protocols. Isolated CD4^+^ T cells were resuspended in RPMI-1640 supplemented with 10% heat-inactivated healthy human type AB serum and Penicillin/Streptomycin (Nacalai Tesque) and used immediately. Cell viability was measured as described above, with a cell concentration set to 4 x 10^4^ cells/well in 96 well culture plates.

### 4.12 Primary CD4^+^ T-cell isolation from HIV patient samples and *in vitro* reactivation

The isolation of primary cells from HIV-positive patients and the *in vitro* reactivation experiments were conducted as described previously (53). In brief, peripheral blood samples were collected from HIV-infected patients who had been receiving combined ART for at least 3 years (Table 1). All patients maintained a low viral load (20 copies/ml) during therapy. CD4^+^ T-cell counts in peripheral blood samples ranged from 324 to 670 cells/mm^3^ (average, 514 cells/mm^3^), and plasma viral loads were 20 copies/ml as measured by qPCR (Roche Applied Science, COBAS AmpliPrep/COBAS TaqMan HIV-1 Test version 2.0) at the time of study enrolment. Written informed consent was obtained from all patients. The Ethics Committee at Kagoshima University and the Japan Institute for Health Security approved this study (NCGM-S-002259-08). This study abided by the principles of the Declaration of Helsinki. Whole PBMCs were separated by density-gradient centrifugation with LymphoprepTM (STEMCELL Technologies), and CD4^+^ T-cells were purified using the MojoSortTM human CD4 T-cell isolation kit (BioLegend) according to the manufacturer’s instructions. Purified CD4^+^ T-cells were plated at a density of 2.5 × 10^6^ cells/ml and treated with 100 nM PMA plus 2 µM ionomycin, 50 µM 2-HSB, 0.2 µM Prostratin, and 1 µM JQ1 for 24 hours. Cells were then harvested and total RNA was extracted using an RNeasy mini kit (Qiagen, Hilden, Germany), following the manufacturer’s protocols. Quantitative real-time PCR analyses for intracellular HIV RNA were carried out using One Step PrimeScript Ⅲ RT-qPCR Mix (TaKaRa Bio). The primer pairs 5’-TGTGTGCCCGTCTGTTGTGT-3’, and 5’-GAGTCCTGCGTCGAGAGAGC-3’ were used for HIV RNA detection. HIV RNA copy numbers were normalized to RNA input. The number of HIV–RNA copies was calculated using a standard curve obtained from serially-diluted HIV plasmids, and normalized values (HIV-RNA copies/input RNA (ng)) with each drug were compared with those without drug; the relative increase in HIV-1 RNA levels in the presence of a drug was then determined.

**Table.** Clinical characteristics of HIV+ participants of this study.

| Participant ID | Sex | Age | VL<sup>a</sup> (copies/mL) | CD4 count<sup>a</sup> (cells/mm<sup>3</sup>) | cART<sup>b</sup> | Duration of therapy (years) |
| --- | --- | --- | --- | --- | --- | --- |
| Participant 1 | M | 39 | <20 | 634 | BIC/TAF/FTC | 6 |
| Participant 2 | M | 56 | <20 | 324 | BIC/TAF/FTC | 20 |
| Participant 3 | M | 50 | <20 | 428 | BIC/TAF/FTC | 5 |
| Participant 4 | M | 45 | <20 | 670 | BIC/TAF/FTC | 3 |

<sup>a</sup>VL and CD4 count were measured at the time of the study.
<sup>b</sup>BIC, bictegravir; DTG, dolutegravir; FTC, emtricitabine; TAF, tenofovir alafenamide fumarate.

### 4.12 Statistical analysis

Statistical analysis was performed using Graphad Prism version 10. Means from at least three independent experiments with error bars indicating standard deviation (SD) are presented in the Results. Statistical significance between means was determined using the unpaired two-tailed Student’s t-test or Analysis of Variance (ANOVA) and Tukey’s post-hoc multiple comparison test. For multiple t-test, Benjamini, Krieger and Yekutieli’s method was used to adjust *P*-values (54). *P*-values <0.05 were considered significant.

## Acknowledgement

This work was supported by: grant 22fk0410052s0101, 23fk0410052h0002, 24fk0410052s0103 for Research Program on HIV/AIDS from Japan Agency for Medical Research and Development to HT; MEXT, the establishment of university fellowships towards the creation of science technology innovation under Grant Number JPMJFS2109 to YH; JST SPRING under Grant Number JPMJSP2120 to YH.

## CRediT authorship contribution statement

**Yuichiro Hara:** Writing – original draft, Writing – review & editing, Methodology, Funding acquisition, Investigation, Formal analysis, Visualisation, Conceptualisation. **Haruki Kitamura**: Investigation, Formal analysis. **Kouki Matsuda:** Writing – original draft, Writing – review & editing, Methodology, Investigation, Resources, Formal analysis. **Sayaka Sukegawa:** Investigation. **Kosuke Tanimoto:** Formal analysis, Visualisation, Software. **Chieko Fujisaki**: Writing – original draft, Writing – review & editing, Methodology, Investigation, Resources, Formal analysis. **Kenji Maeda:** Writing – original draft, Writing – review & editing, Resources, Supervision. **Hiroaki Takeuchi:** Writing – review & editing, Project administration, Supervision, Funding acquisition, Resources, Methodology, Conceptualisation.

---

**Supplementary Fig. 1.**
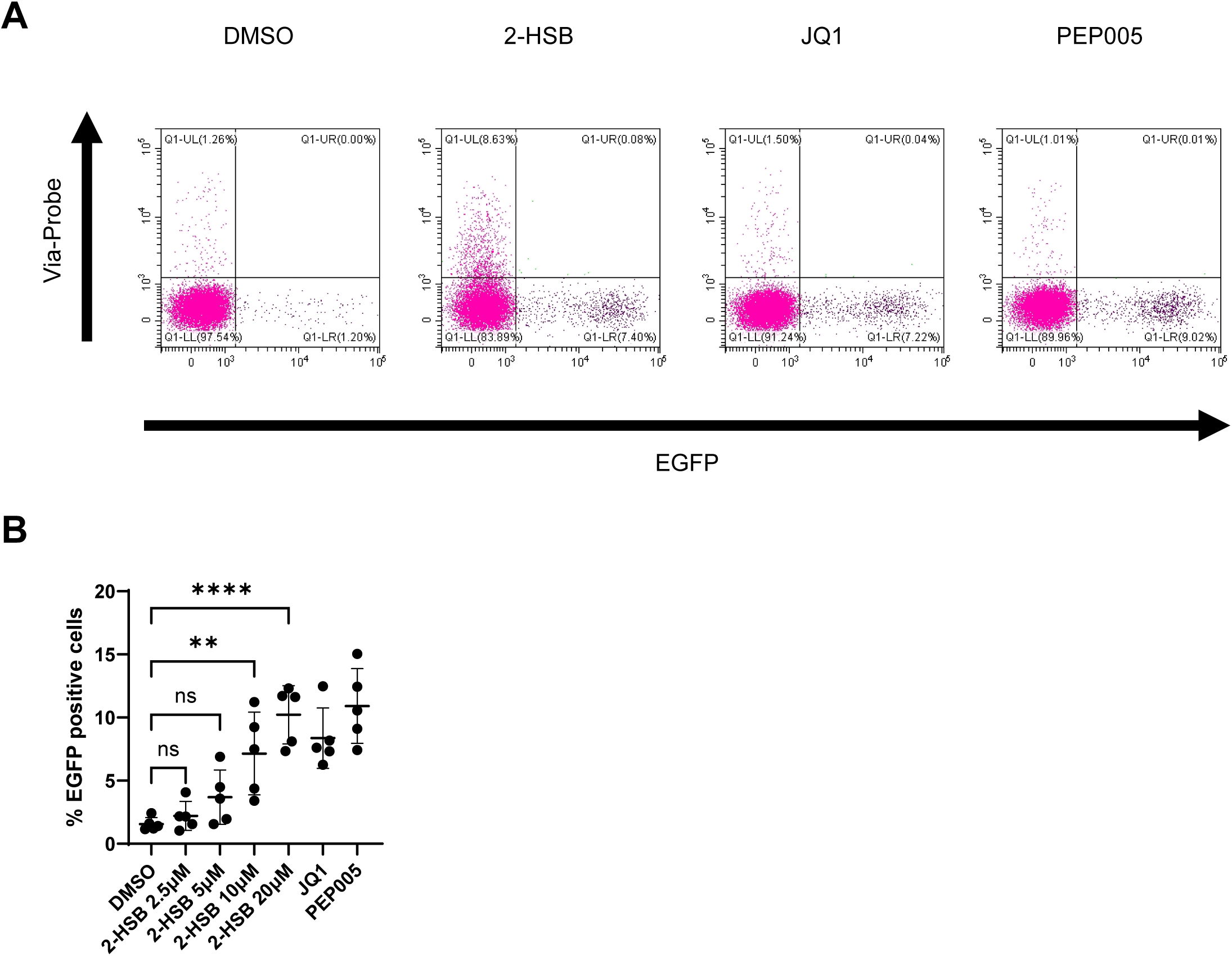
Flow cytometry of J-Lat 10.6 cells treated with LRAs. (A) Flow cytometry panel of 10.6 cells treated with DMSO, 2-HSB, JQ1 or PEP005 for 24 hours. The X-axis shows the FITC channel (GFP) and the Y-axis shows the PC5.5 channel (Via-Probe). (B) The absolute percentage of GFP-positive cells in 10.6 cells after treatment with various compounds. The fold-change relative to DMSO-treated cells is summarized in Figure 1C.

**Supplementary Fig. 2.**
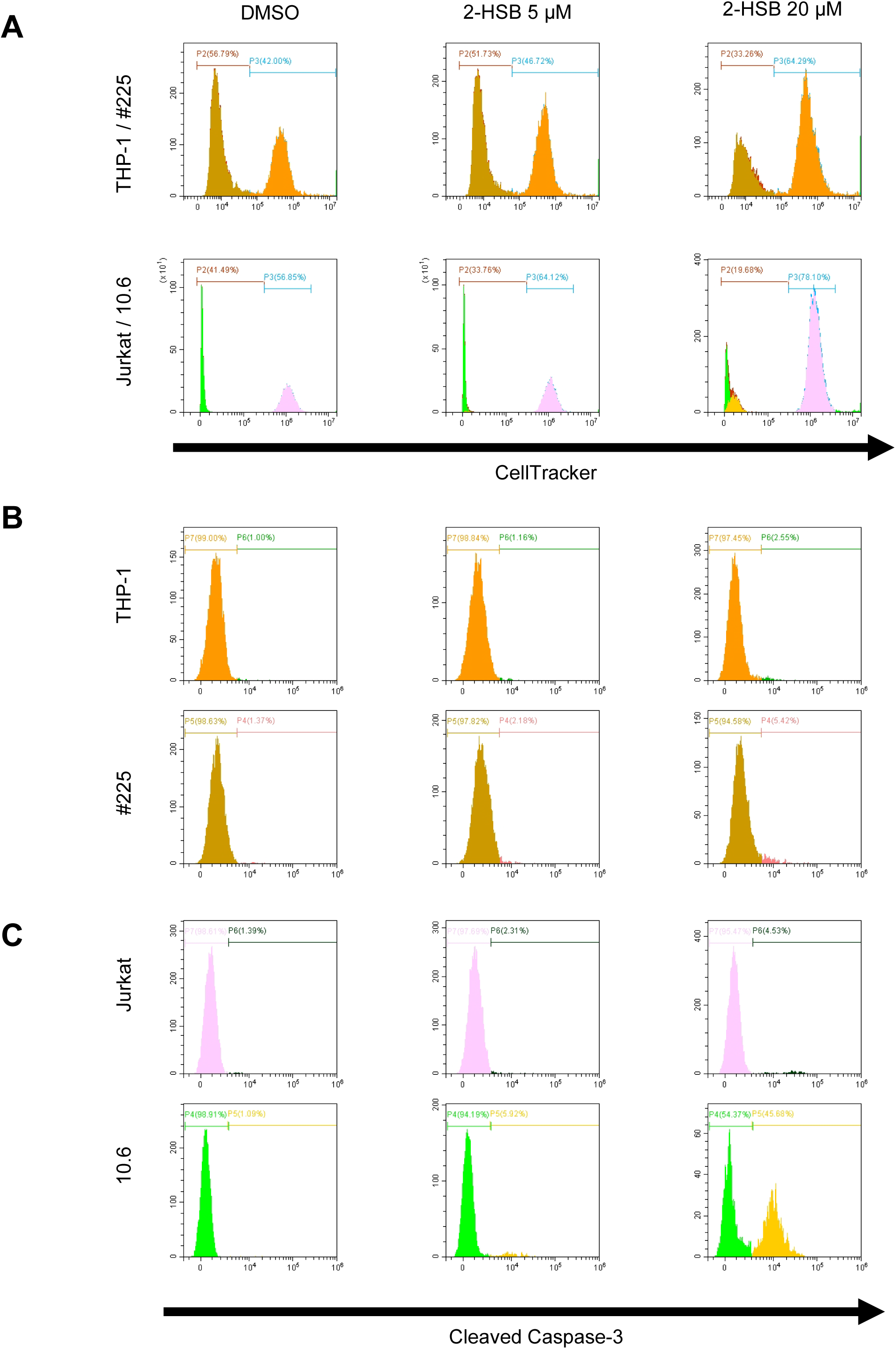
Flow cytometric analysis of selective cytotoxicity in mixed cell populations treated with 2-HSB. Mixed populations of labeled (HIV-1-negative) and unlabeled (HIV-1-positive) cells were treated with DMSO or 2-HSB (5 or 20 μM) for 24 hours. (A) Representative histograms showing the population distribution of labeled vs. unlabeled cells. (Upper panels) Co-culture of THP-1 (labeled) and #225 (unlabeled). (Lower panels) Co-culture of Jurkat (labeled) and 10.6 (unlabeled). (B–C) Histograms displaying the percentage of cleaved Caspase-3-positive cells within each population defined in (A). (B) Analysis of monocytic cells: THP-1 (upper) and #225 (lower). (C) Analysis of T-lymphocytic cells: Jurkat (upper) and 10.6 (lower). Representative data from three independent experiments are shown.

**Supplementary Fig. 3.**
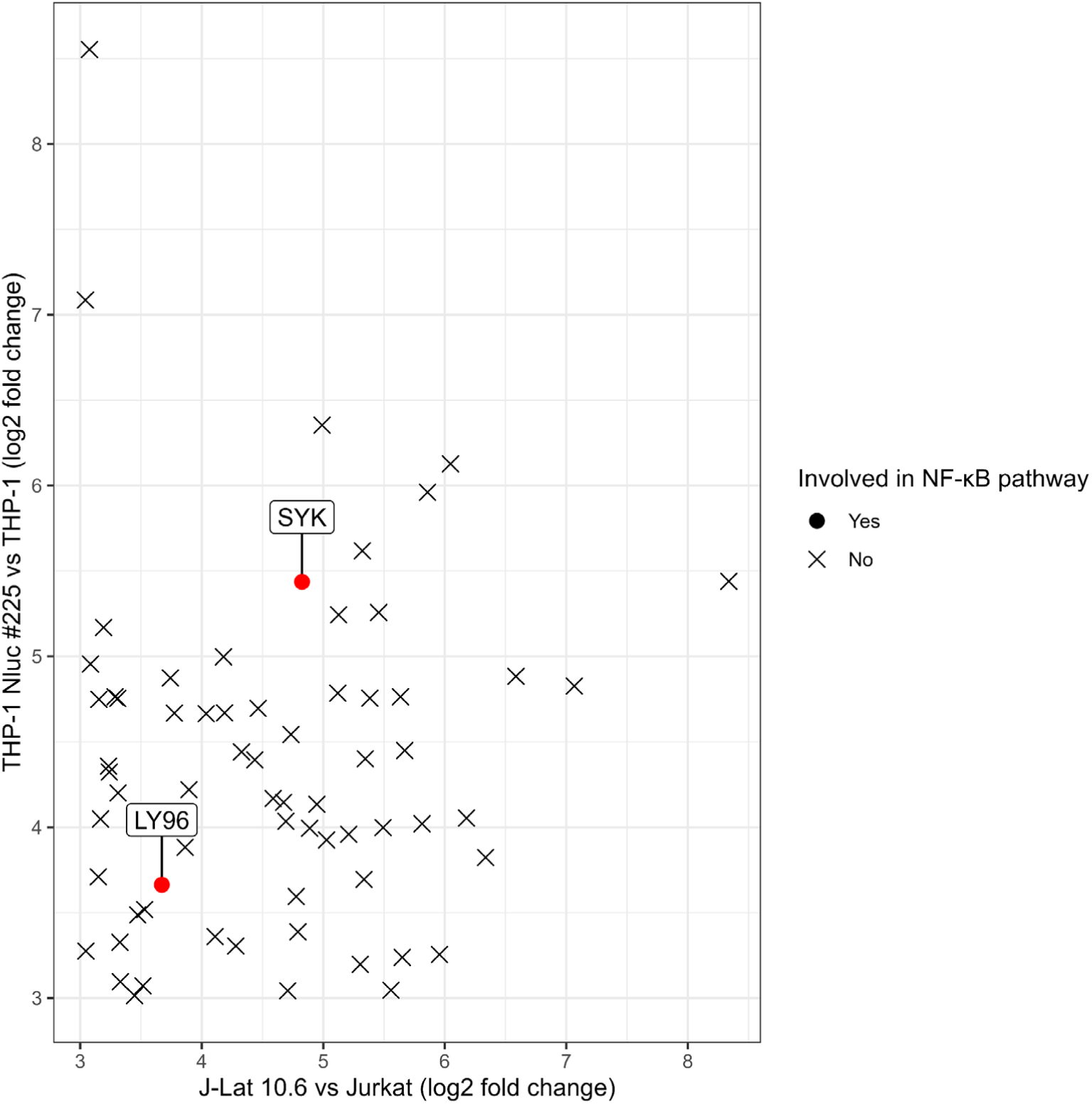
Microarray profiling of genes upregulated in provirus-positive cells. THP-1, THP-1 Nluc #225, Jurkat and J-Lat 10.6 cells were treated with 10 µM 2-HSB for 24 hours. Differential expression analysis was performed to identify genes upregulated exclusively in provirus-positive cells relative to their respective parental lines (THP-1 or Jurkat). Genes exhibiting a log2 fold change > 3 (> 8-fold increase) are plotted. Genes involved in the NF-κB pathway are highlighted in red. Data represent a single exploratory screening experiment (n = 1).

**Supplementary Table 1.**
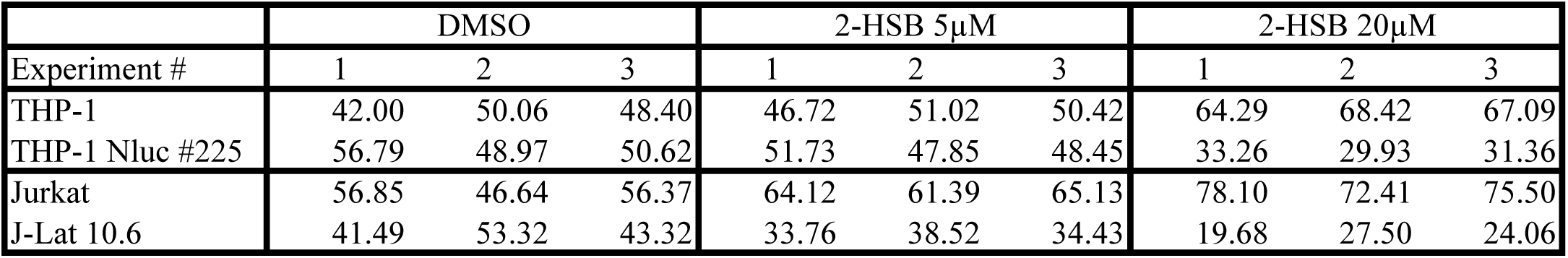
Population size of cell lines after treatment by 2-HSB.

**Supplementary Table 2.**
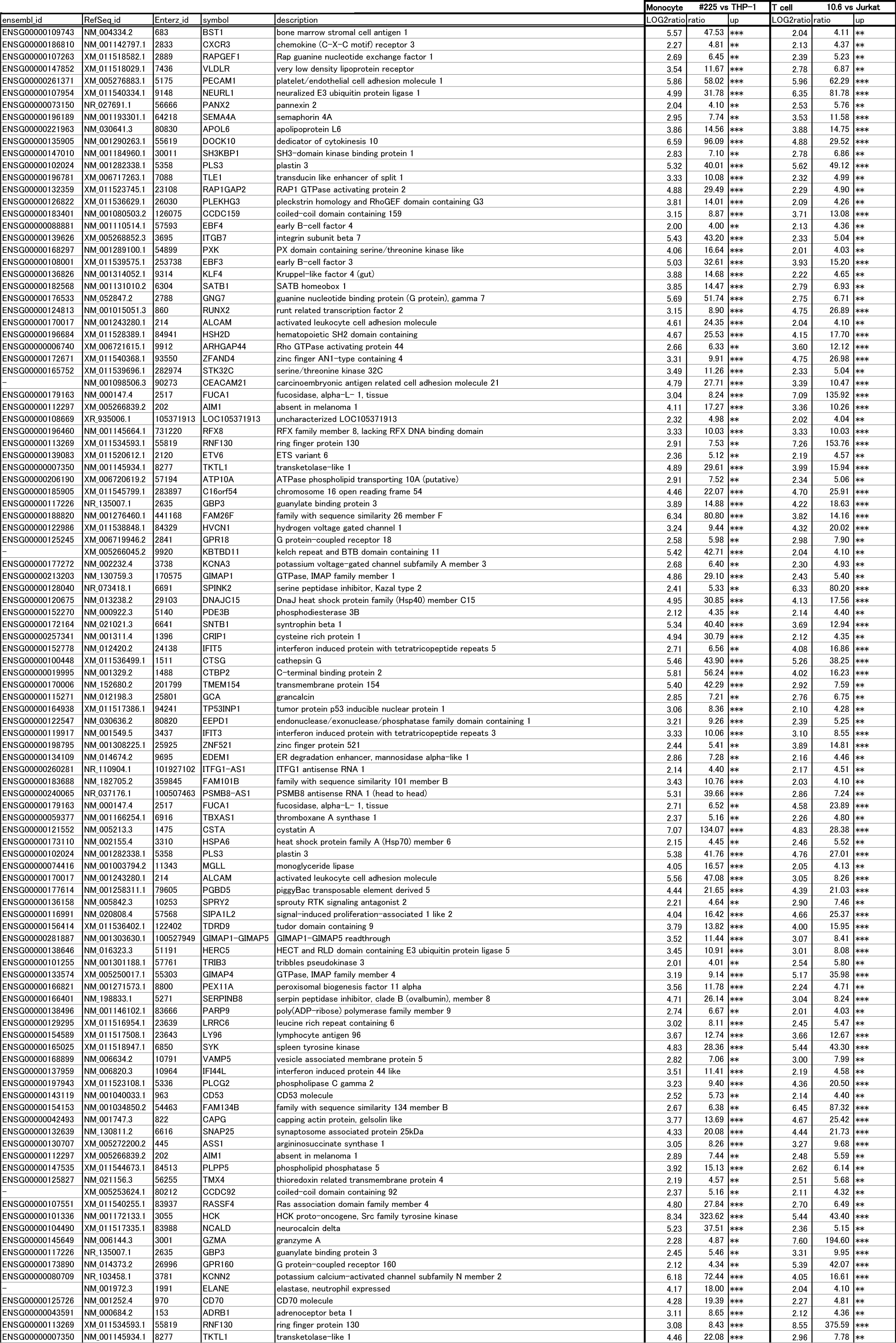
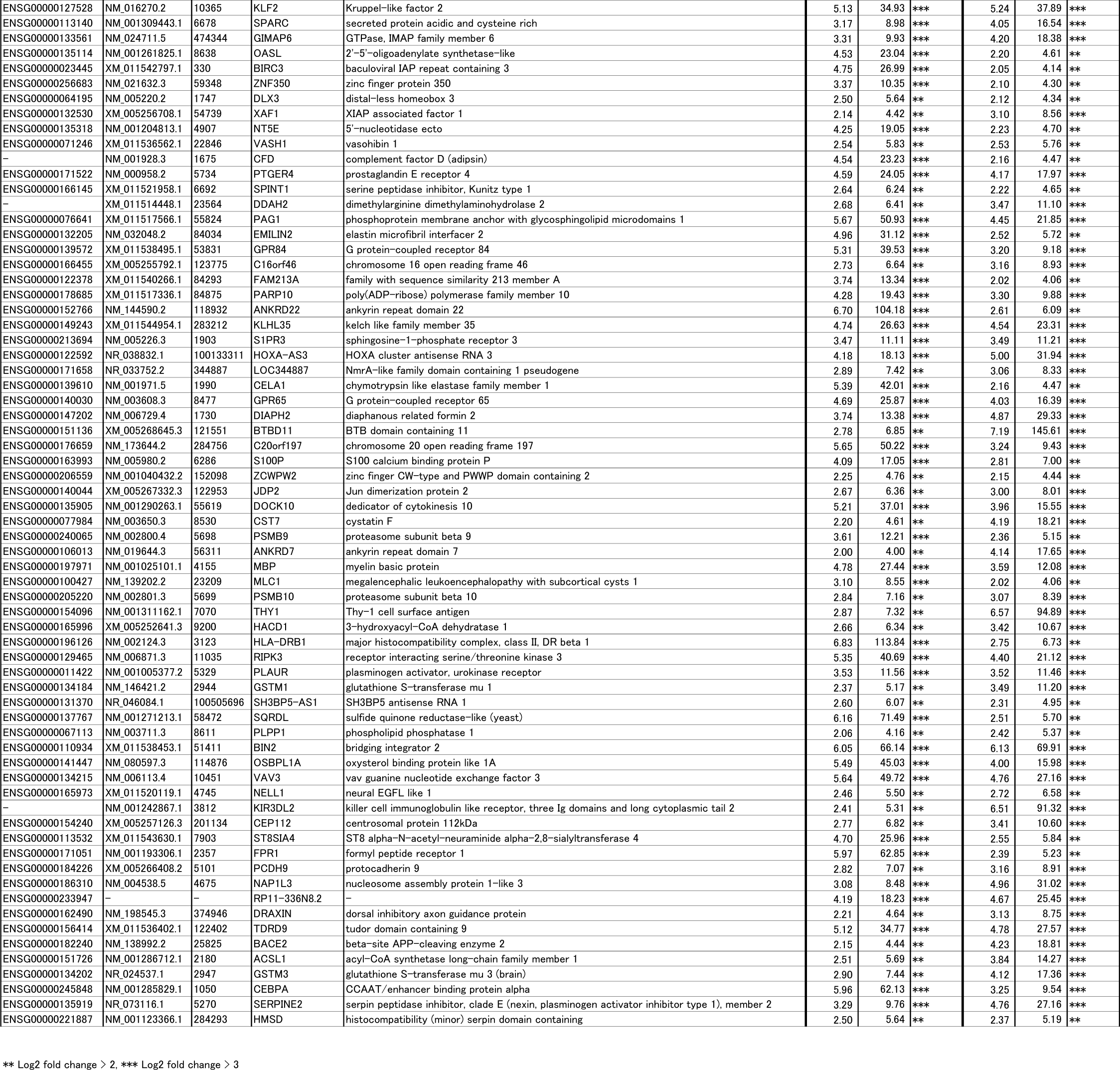
List of genes identified in microarray transcriptome analysis.

## References

1. Mitsuya H, Weinhold KJ, Furman PA, St Clair MH, Lehrman SN, Gallo RC, et al. 3’-Azido-3’-deoxythymidine (BW A509U): an antiviral agent that inhibits the infectivity and cytopathic effect of human T-lymphotropic virus type III/lymphadenopathy-associated virus in vitro. Proceedings of the National Academy of Sciences. 1985;82(20):7096–100.

2. Palmisano L, Vella S. A brief history of antiretroviral therapy of HIV infection: success and challenges. Ann Ist Super Sanita. 2011;47(1):44–8.

3. Hütter G, Nowak D, Mossner M, Ganepola S, Müßig A, Allers K, et al. Long-Term Control of HIV by *CCR5* Delta32/Delta32 Stem-Cell Transplantation. New England Journal of Medicine. 2009;360(7):692–8.

4. Davenport MP, Khoury DS, Cromer D, Lewin SR, Kelleher AD, Kent SJ. Functional cure of HIV: the scale of the challenge. Nat Rev Immunol. 2019;19(1):45–54.

5. Castro-Gonzalez S, Colomer-Lluch M, Serra-Moreno R. Barriers for HIV Cure: The Latent Reservoir. AIDS Res Hum Retroviruses. 2018;34(9):739–59.

6. Xiao Q, Guo D, Chen S. Application of CRISPR/Cas9-Based Gene Editing in HIV-1/AIDS Therapy. Front Cell Infect Microbiol. 2019;9:69.

7. Koup RA, Douek DC. Vaccine design for CD8 T lymphocyte responses. Cold Spring Harb Perspect Med. 2011;1(1):a007252.

8. Blazkova J, Chun TW, Belay BW, Murray D, Justement JS, Funk EK, et al. Effect of histone deacetylase inhibitors on HIV production in latently infected, resting CD4(+) T cells from infected individuals receiving effective antiretroviral therapy. J Infect Dis. 2012;206(5):765–9.

9. Lopes JR, Chiba DE, Dos Santos JL. HIV latency reversal agents: A potential path for functional cure? Eur J Med Chem. 2021;213:113213.

10. Debrabander Q, Hensley KS, Psomas CK, Bramer W, Mahmoudi T, van Welzen BJ, et al. The efficacy and tolerability of latency-reversing agents in reactivating the HIV-1 reservoir in clinical studies: a systematic review. J Virus Erad. 2023;9(3):100342.

11. Fidler S, Stohr W, Pace M, Dorrell L, Lever A, Pett S, et al. Antiretroviral therapy alone versus antiretroviral therapy with a kick and kill approach, on measures of the HIV reservoir in participants with recent HIV infection (the RIVER trial): a phase 2, randomised trial. Lancet. 2020;395(10227):888–98.

12. Kitamura H, Sukegawa S, Matsuda K, Tanimoto K, Kobayakawa T, Takahashi K, et al. 4-phenylquinoline-8-amine induces HIV-1 reactivation and apoptosis in latently HIV-1 infected cells. Biochem Biophys Res Commun. 2023;641:139–47.

13. Gohda J, Suzuki K, Liu K, Xie X, Takeuchi H, Inoue JI, et al. BI-2536 and BI-6727, dual Polo-like kinase/bromodomain inhibitors, effectively reactivate latent HIV-1. Sci Rep. 2018;8(1):3521.

14. Jordan A, Bisgrove D, Verdin E. HIV reproducibly establishes a latent infection after acute infection of T cells *in vitro*. The EMBO Journal. 2003;22(8):1868–77.

15. Brogdon J, Ziani W, Wang X, Veazey RS, Xu H. In vitro effects of the small-molecule protein kinase C agonists on HIV latency reactivation. Sci Rep. 2016;6:39032.

16. Li Z, Guo J, Wu Y, Zhou Q. The BET bromodomain inhibitor JQ1 activates HIV latency through antagonizing Brd4 inhibition of Tat-transactivation. Nucleic Acids Res. 2013;41(1):277–87.

17. Abner E, Jordan A. HIV “shock and kill” therapy: In need of revision. Antiviral Res. 2019;166:19–34.

18. Kim Y, Anderson JL, Lewin SR. Getting the “Kill” into “Shock and Kill": Strategies to Eliminate Latent HIV. Cell Host Microbe. 2018;23(1):14–26.

19. Porter AG, Jänicke RU. Emerging roles of caspase-3 in apoptosis. Cell Death & Differentiation. 1999;6(2):99–104.

20. DeLuca C, Kwon H, Lin R, Wainberg M, Hiscott J. NF- κ B activation and HIV-1 induced apoptosis. Cytokine & Growth Factor Reviews. 1999;10(3):235–53.

21. Takada Y, Aggarwal BB. TNF activates Syk protein tyrosine kinase leading to TNF-induced MAPK activation, NF-kappaB activation, and apoptosis. J Immunol. 2004;173(2):1066–77.

22. Takano T, Sada K, Yamamura H. Role of Protein-Tyrosine Kinase Syk in Oxidative Stress Signaling in B Cells. Antioxidants & Redox Signaling. 2002;4(3):533–41.

23. Rochat MA, Schlaepfer E, Speck RF. Promising Role of Toll-Like Receptor 8 Agonist in Concert with Prostratin for Activation of Silent HIV. J Virol. 2017;91(4).

24. Hisamichi H, Naito R, Toyoshima A, Kawano N, Ichikawa A, Orita A, et al. Synthetic studies on novel Syk inhibitors. Part 1: Synthesis and structure-activity relationships of pyrimidine-5-carboxamide derivatives. Bioorg Med Chem. 2005;13(16):4936–51.

25. Zack JA, Arrigo SJ, Weitsman SR, Go AS, Haislip A, Chen IS. HIV-1 entry into quiescent primary lymphocytes: molecular analysis reveals a labile, latent viral structure. Cell. 1990;61(2):213–22.

26. Kandathil AJ, Sugawara S, Balagopal A. Are T cells the only HIV-1 reservoir? Retrovirology. 2016;13(1):86.

27. Ruelas DS, Greene WC. An integrated overview of HIV-1 latency. Cell. 2013;155(3):519–29.

28. Cummins NW, Badley AD. Mechanisms of HIV-associated lymphocyte apoptosis: 2010. Cell Death Dis. 2010;1(11):e99.

29. Bartz Steven R, Emerman M. Human Immunodeficiency Virus Type 1 Tat Induces Apoptosis and Increases Sensitivity to Apoptotic Signals by Up-Regulating FLICE/Caspase-8. Journal of Virology. 1999;73(3):1956–63.

30. Ali A, Mishra R, Kaur H, Chandra Banerjea A. HIV-1 Tat: An update on transcriptional and non-transcriptional functions. Biochimie. 2021;190:24–35.

31. Shirakawa K, Chavez L, Hakre S, Calvanese V, Verdin E. Reactivation of latent HIV by histone deacetylase inhibitors. Trends Microbiol. 2013;21(6):277–85.

32. Jiang G, Dandekar S. Targeting NF-kappaB signaling with protein kinase C agonists as an emerging strategy for combating HIV latency. AIDS Res Hum Retroviruses. 2015;31(1):4–12.

33. Kulkosky J, Culnan DM, Roman J, Dornadula G, Schnell M, Boyd MR, et al. Prostratin: activation of latent HIV-1 expression suggests a potential inductive adjuvant therapy for HAART. Blood. 2001;98(10):3006–15.

34. Alimonti JB, Ball TB, Fowke KR. Mechanisms of CD4+ T lymphocyte cell death in human immunodeficiency virus infection and AIDS. J Gen Virol. 2003;84(Pt 7):1649–61.

35. Khan SZ, Hand N, Zeichner SL. Apoptosis-induced activation of HIV-1 in latently infected cell lines. Retrovirology. 2015;12:42.

36. Palmer S, Josefsson L, Coffin JM. HIV reservoirs and the possibility of a cure for HIV infection. J Intern Med. 2011;270(6):550–60.

37. Wong ME, Jaworowski A, Hearps AC. The HIV Reservoir in Monocytes and Macrophages. Front Immunol. 2019;10:1435.

38. Sacha JB, Ndhlovu LC. Strategies to target non-T-cell HIV reservoirs. Curr Opin HIV AIDS. 2016;11(4):376–82.

39. Rice AP. The HIV-1 Tat Protein: Mechanism of Action and Target for HIV-1 Cure Strategies. Curr Pharm Des. 2017;23(28):4098–102.

40. Huang L, Bosch I, Hofmann W, Sodroski J, Pardee Arthur B. Tat Protein Induces Human Immunodeficiency Virus Type 1 (HIV-1) Coreceptors and Promotes Infection with both Macrophage-Tropic and T-Lymphotropic HIV-1 Strains. Journal of Virology. 1998;72(11):8952–60.

41. Luznik L, Kraus G, Guatelli J, Richman D, Wong-Staal F. Tat-independent replication of human immunodeficiency viruses. J Clin Invest. 1995;95(1):328–32.

42. Emiliani S, Fischle W, Ott M, Van Lint C, Amella Carol A, Verdin E. Mutations in the tat Gene Are Responsible for Human Immunodeficiency Virus Type 1 Postintegration Latency in the U1 Cell Line. Journal of Virology. 1998;72(2):1666–70.

43. Badley AD, Pilon AA, Landay A, Lynch DH. Mechanisms of HIV-associated lymphocyte apoptosis. Blood. 2000;96(9):2951–64.

44. Appay V, Sauce D. Immune activation and inflammation in HIV-1 infection: causes and consequences. J Pathol. 2008;214(2):231–41.

45. Van Opdenbosch N, Lamkanfi M. Caspases in Cell Death, Inflammation, and Disease. Immunity. 2019;50(6):1352–64.

46. Brady T, Agosto LM, Malani N, Berry CC, O’Doherty U, Bushman F. HIV integration site distributions in resting and activated CD4+ T cells infected in culture. AIDS. 2009;23(12):1461–71.

47. Maldarelli F, Wu X, Su L, Simonetti FR, Shao W, Hill S, et al. Specific HIV integration sites are linked to clonal expansion and persistence of infected cells. Science. 2014;345(6193):179–83.

48. Matsuda Y, Kobayashi-Ishihara M, Fujikawa D, Ishida T, Watanabe T, Yamagishi M. Epigenetic heterogeneity in HIV-1 latency establishment. Sci Rep. 2015;5:7701.

49. Dobin A, Davis CA, Schlesinger F, Drenkow J, Zaleski C, Jha S, et al. STAR: ultrafast universal RNA-seq aligner. Bioinformatics. 2013;29(1):15–21.

50. Li B, Dewey CN. RSEM: accurate transcript quantification from RNA-Seq data with or without a reference genome. BMC Bioinformatics. 2011;12(1):323.

51. Love MI, Huber W, Anders S. Moderated estimation of fold change and dispersion for RNA-seq data with DESeq2. Genome Biol. 2014;15(12):550.

52. Gu Z, Eils R, Schlesner M. Complex heatmaps reveal patterns and correlations in multidimensional genomic data. Bioinformatics. 2016;32(18):2847–9.

53. Matsuda K, Kobayakawa T, Tsuchiya K, Hattori SI, Nomura W, Gatanaga H, et al. Benzolactam-related compounds promote apoptosis of HIV-infected human cells via protein kinase C-induced HIV latency reversal. J Biol Chem. 2019;294(1):116–29.

54. Benjamini Y, Krieger AM, Yekutieli D. Adaptive Linear Step-up Procedures That Control the False Discovery Rate. Biometrika. 2006;93(3):491–507.

